# Investigation of refractive index dynamics during embryo development using digital holographic microscopy

**DOI:** 10.1101/2023.04.17.537152

**Authors:** George O. Dwapanyin, Darren J. X. Chow, Tiffany C. Y. Tan, Nicolas S. Dubost, Josephine M. Morizet, Kylie R. Dunning, Kishan Dholakia

**Author notes:** These authors contributed equally (George O. Dwapanyin, Darren J. X. Chow).

## Abstract

Embryo quality is a crucial factor affecting live birth outcomes. However, an accurate diagnostic for embryo quality remains elusive in the *in vitro* fertilization clinic. Determining physical parameters of the embryo may offer key information for this purpose. Here, we demonstrate that digital holographic microscopy (DHM) can rapidly and non-invasively assess the refractive index of mouse embryos. We showed that DHM can detect spatio-temporal changes in refractive index during embryo development that are reflective of its lipid content. As accumulation of intracellular lipid is known to compromise embryo health, DHM may prove beneficial in developing an accurate, non-invasive, multimodal diagnostic.

## 1. Introduction

Pre-implantation embryo development describes events that occur following the fusion of a sperm and an oocyte through to the blastocyst-stage of embryo development. It is at this stage that the embryo is capable of implanting into the uterus and establishing a pregnancy. The quality, or developmental potential, of an embryo is critical as it dictates downstream pregnancy success and ultimately, the birth of a live offspring.

Within an *in vitro* fertilization (IVF) clinic, embryo quality is routinely assessed by visual inspection (i.e. is the embryo developing in a time-appropriate manner) and via an invasive biopsy (to assess DNA content of the biopsied sample). However, these approaches have failed to improve the success rate of IVF, which has remained stagnant for more than a decade: ∼ 30% of initiated cycles result in a live birth [1]. The inability of these assessments to improve IVF success is likely due to their subjective nature or the diagnostic approach causing inaccuracies [2]. Furthermore, biopsy requires the removal of a subset of cells from the embryo which is both highly invasive and may increase the likelihood of a pathology developing during pregnancy post-embryo transfer [3].

Towards developing an objective and accurate diagnostic for embryo quality, there has been a recent surge in the use of optical approaches within the field. For example, identification of chromosomal aberrations, traditionally performed via embryo biopsy - may now be determined using hyperspectral imaging [4] or fluorescence lifetime imaging microscopy (FLIM) [5]. Further, optical coherence microscopy (OCM), a label-free and non-invasive imaging technique, has been shown to detect changes in subcellular architecture that were associated with embryo quality [6, 7]. These optical approaches obviate the need for invasive removal of cells from the embryo or subjective visual inspection by directly accessing key information within an embryo, such as metabolic activity. Separately, biodynamic imaging (BDI), based on low-coherence dynamic light scattering and low-coherence digital holography has also been used to assess the viability and metabolic activity of oocytes and embryos [8, 9]. However, BDI relies on the detection of subcellular motion within the living tissue to create an endogenous imaging contrast and not the physical parameters of the cell itself. To date, little attention has been paid to the physical parameters of the embryo, including dynamic changes in refractive properties during development, which may be indicative of embryo quality. Lipids are established to have a higher refractive index (RI) than protein [10]. Convincingly, in microalgae, changes in RI have been shown to correlate with intracellular lipid content [11]. In the embryo, lipids are a potent source of energy [12–16], and dysregulation of their metabolism (i.e. increased intracellular lipid) is associated with impaired embryo quality [17]. Collectively, these studies indicate the value of observing RI as a potential indicator of embryo health.

In the last decade, digital holographic microscopy (DHM) has emerged as a powerful method to extract morphological information from biological samples [18, 19]. In particular, DHM enables quantitative phase imaging that can lead to the determination of RI in a spatial manner. We note that in previous work, Wacogne et al [20] measured the transmission spectra and used the lensing action of an oocyte to determine its refractive index. Importantly, this approach fails to provide spatial information on RI but instead reports a value for the whole sample, which may exclude developmentally important information. The study also reported very high values of refractive index for the oocyte (average RI = 1.8), well beyond what might be expected in biological cells.

In this paper, we show the first application of DHM on mouse pre-implantation embryos to study dynamic physical changes of RI - in a spatio-temporal manner. We characterized RI across pre-implantation embryo development and determined whether changes in RI were associated with embryo quality. To the best of our knowledge, this is the first instance where RI has been linked to embryo quality. As such, our study shows the potential for DHM, a rapid and label-free optical approach, to objectively play a role in the assessment of embryo quality.

## 2. Materials and methods

### 2.1. DHM setup

The DHM imaging system is shown in Figure 1. Its design is based on an off-axis Mach-Zehnder transmission interferometric configuration with a single-frequency free-space continuous wave laser (Coherent Sapphire SF NX, 532 nm, 75 mW). Vertical and horizontal polarization components of the laser output were separated using a polarizing beam splitter (PBS) (Thorlabs, CCM1-PBS251/M). The vertical polarization was coupled into a 50:50 single-mode fibre beam splitter (FBS) (Thorlabs, FC532-50B-FC) using a 10X objective (Thorlabs, RMS10X) while the horizontal component was dumped into a beam block (BB). The beam from one arm was coupled to the object path and collimated with a fibre collimator (Thorlabs, F220FC-A). It was then expanded with a lens (L1) before being focused through a 10X objective (Obj.1) (Mitutoyo UK, M Plan Apo 10X 0.23NA, WD 34.0 mm) placed along the same path as the bright field illumination. This beam expansion was performed to fill the back aperture of the objective. The object beam passed through the sample placed in a glass bottom petri dish and positioned on a stage heated to 37 °C via a peltier heater (Digi-Key electronics). Collection was performed with a 40X objective (Obj.2) (Nikon 0.65 NA). Images were focused on a charge-coupled device (CCD) camera (Ximea XiQ MQ013MG-E2) with a 200 mm tube lens with the images having a transverse resolution of 1 μm.

**Fig. 1.**
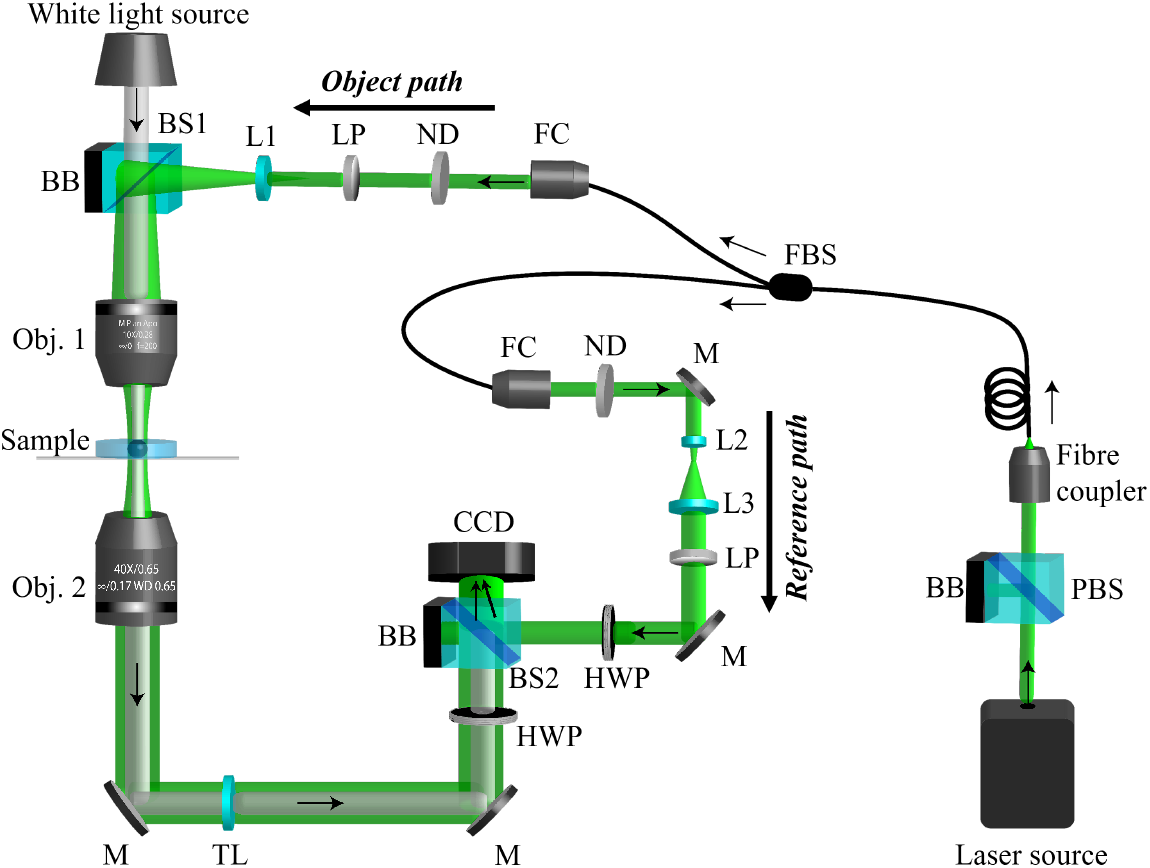
Schematic diagram of the DHM system. The system is based on a Mach-Zehnder interferometer. Vertically polarized light from the laser is split into an object and a reference beam path. The object beam illuminates the sample before recombining with the reference beam to form a hologram. The off-axis geometry for interference is achieved by a beam splitter placed in front of the CCD camera. The system has a magnification of 40X and a 154 × 192 μm^2^ maximum field of view. Arrows indicate beam direction. The abbreviations refer to: PBS: polarization beam splitter (Thorlabs, CCM1-PBS251/M), BB: beam block, FBS: Fibre beam splitter (Thorlabs, FC532-50B-FC), FC: fibre collimator (Thorlabs, F220FC-A), ND: Neutral density wheel, LP: Linear polarizer (LPVISE100-A), lenses L1=100 mm, L2=25 mm and L3=75 mm. BS: Non-polarizing beam splitter (Thorlabs, BS016), Obj. 1: Mitutoyo Plan Apochromat (10X, 0.28 NA, 34 mm WD) objective, Obj. 2: Nikon CFI (40X, 0.65 NA) objective, M: Mirror, TL: 200 mm tube lens, HWP: Half waveplate (WPH10M-532), CCD: Charge-coupled device camera (Ximea XiQ MQ013MG-E2).

The beam from the reference arm was collimated and expanded through a 3X magnification telescope formed by lenses L2 and L3. The object and reference paths were recombined using a 50:50 non-polarizing beam splitter (BS2) (Thorlabs, BS016) at an angle to create an off-axis hologram on the CCD camera. This camera was set to accumulate 16-bit images with a frame rate of 60 fps and an exposure time of 33.3 ms. The system has a 154 × 192 μm^2^ maximum field of view. The power in both arms was controlled with round variable neutral density wheels (ND) (Thorlabs, NDC-50C-2M-A) with powers of 30 μW and 25 μW in the object and reference arm respectively. A combination of linear polarizers (LP) (Thorlabs LPVISE100-A) and zero-order half waveplates (HWP) (Thorlabs-WPH10M-532) in both arms ensured equivalent linearly polarized beams that could be interfered to form the hologram on the CCD. White light source for brightfield images was provided by a broad mounted LED source (Thorlabs MBB1L3) mounted on top of the signal arm.

### 2.2. Culture media preparation

All embryo culture took place in media overlaid with paraffin oil (Merck Group, Darmstadt, Germany) at 37°C in a humified incubator set at 5% O_2_ and 6% CO_2_ balanced in N_2_. Culture dishes were pre-equilibrated for at least 4 h prior to use. All handling procedures were performed on microscopes fitted with heating stages calibrated to maintain media in dishes at 37°C. All culture media were supplemented with 4 mg/ml low fatty acid bovine serum albumin (BSA, MP Biomedicals, AlbumiNZ, Auckland, NZ) unless specified otherwise. Oviducts were collected in filtered Research Wash medium (ART Lab Solutions, SA, Australia) and embryos were cultured in filtered Research Cleave medium (ART Lab Solutions, SA, Australia).

### 2.3. Collection of embryos

Female (21-23 days) CBA x C57BL/6 first filial (CBAF1) generation mice were obtained from Laboratory Animal Services (University of Adelaide, Australia) and maintained on a 12 h light: 12 h dark cycle with rodent chow and water provided *ad libitum*. All studies were approved by both University of St. Andrews’ School of Biology Ethics Committee (SEC20001) and the University of Adelaide’s Animal Ethics Committee (M-2019-097) and were conducted in accordance with the Australian Code of Practice for the Care and Use of Animals for Scientific Purposes. Female mice were administered intraperitoneally (i.p.) with 5 IU of equine chorionic gonadotropin (eCG; Folligon, Braeside, VIC, Australia), followed by 5 IU human chorionic gonadotrophin (hCG, i.p.; Kilsyth, VIC, Australia) 46 h later. Female mice were then mated overnight with male mice of proven fertility. At 47 h post-hCG, females were culled by cervical dislocation and the oviducts dissected to isolate 2-cell embryos. Two-cell stage embryos were released from the oviducts by gently flushing the oviduct using pre-warmed Research Wash medium supplemented with 4 mg/ml BSA using a 29-gauge insulin syringe with needle (Terumo Australia Pty Ltd, Australia) and subsequently underwent vitrification.

#### 2.3.1. Embryo vitrification and warming

Media used for embryo vitrification and warming were as described in Tan et al [4]. Briefly, 2-cell stage embryos were vitrified with the Cryologic vitrification method, consisting of timely, sequential washes in handling medium followed by 3 min in equilibration solution, and 30 s in vitrification solution, prior to loading onto a Fibreplug straw for storage in liquid nitrogen. For embryo warming, Fibreplugs containing embryos were removed from liquid nitrogen and quickly submerged into a handling medium supplemented with 0.3 M sucrose, followed by sequential washes in handling media with decreasing concentrations of sucrose (0.25, 0.15, and 0 M) for 5 min each. The post-warming survival rate was above 85% for all groups (data not shown). Warmed 2-cell stage embryos were then cultured in Research Cleave medium supplemented with 4 mg/ml BSA either in the absence (*low-lipid* group) or presence of 10% fetal bovine serum (*high-lipid* group) and allowed to develop up until the blastocyst-stage in these conditions. The presence of 10% serum during culture is known to promote embryo developmental rate in bovine [21], however this is at the cost of altering molecular mechanisms that may lead to long-term developmental anomalies after implantation, such as fetal overgrowth. Therefore, we contend that the addition of 10% fetal bovine serum during embryo culture is sufficient to elevate the levels of intracellular lipids within the embryo.

### 2.4. Sample preparation and DHM imaging

For imaging, 2-, 4-, 8-cell, morula and blastocyst-stage embryos for each treatment group were collected at 6-, 10-, 24-, 30-, and 48-h post-warming respectively, and were transferred into pre-warmed 20 μl drops of Research Wash medium supplemented with 4 mg/ml BSA in a glass-bottom imaging dish overlaid with paraffin oil. The Research Wash medium (with a refractive index of 1.3355) provides a physiologically pH-buffered medium for live embryo imaging. To limit thermal shock during transportation between the culture incubator and the DHM imaging system, an enclosed chamber with controlled heating pads was used to provide a constant incubation temperature at 37°C. Bright-field images of embryos were taken using a standard white light source integrated into the imaging system. The total number of embryos imaged at each developmental stage is shown in Supplementary Table S1.

#### 2.4.1. BODIPY 493/503 staining and imaging

Embryos were fixed in 4% paraformaldehyde-phosphate buffer saline for 30 min and rinsed thoroughly in 0.3 mg/mL polyvinyl alcohol in phosphate buffer saline (PBV). Following 1 h incubation at room temperature with BODIPY 493/503 (1 μg/mL; ThermoFisher), embryos were thoroughly washed in PBV and mounted on glass microscope slide and enclosed with a coverslip and a spacer (ThermoFisher Scientific, Waltham, MA, USA) in PBV before proceeding to imaging and analysis. BODIPY 493/503-stained embryos were captured on an Olympus FluoView FV10i confocal laser scanning microscope (Olympus, Tokyo, Japan). Images were acquired at 60X magnification with a water-immersion compatible objective (Olympus, 1.2 NA). Images were captured at 2 μm intervals through the entire embryo and a final z-stack maximum intensity projection was generated. Samples were excited at 488 nm (emission detection wavelength: 490–525 nm) to detect BODIPY-stained cells. Lipid abundance was quantified by fluorescence intensity of BODIPY staining in a z-stack maximum intensity projection generated for each embryo using ImageJ for Windows 10 (Fiji, MD, USA).

### 2.5. Theory and data analysis

DHM is an interferometric phase imaging technique that measures the phase shift of light traversing through transparent samples. For light passing through a sample of varying refractive index, *n*, through a thickness *h* in the *z*−direction, the phase shift *ϕ* encountered by the light can be written as:

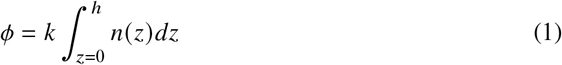

where *k* is the wavenumber given by 2*π*/*λ* with *λ* being the excitation wavelength. For cells of refractive index *n*_*s*_ in a fluid medium on index *n*_*m*_, the phase difference Δ*ϕ* encountered by the light during propagation can be summarized as:

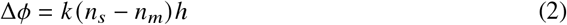

Thus with a knowledge of Δ*ϕ* and *n*_*m*_, the height/thickness *h* and/or the refractive index of the cell *n*_*s*_ can be determined. To determine the thickness at each camera pixel, consider Fig.2. For each *i*^*th*^ pixel, the radius (*r*_*i*(*x,y*)_) from the centre of the circular region of interest (ROI) is given by:

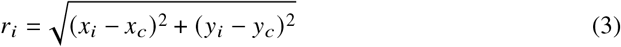

where *x*_*i*_ and *y*_*i*_ represent the coordinates of the *i*^*th*^ pixel while *x*_*c*_ and *y*_*c*_ represent the centre coordinates in X and Y directions. The height, *h*, defined as the distance travelled by a light ray going through the sample at the *i*^*th*^ pixel position thus relates to the radius by the expression:

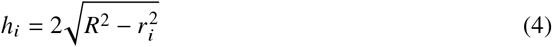

where *R* is the radius from the hemisphere and the factor 2 accounts for the other half of the hemisphere since embryos are known to be near spherical in nature [6]. Substituting Eq. 4 into Eq. 2 and rearranging, the refractive index of the sample at each pixel point can be deduced such that:

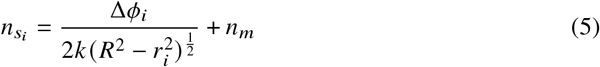

The phase shift Δ*ϕ* is encoded or modulated into the interferogram *I*_*C*_ detected by the CCD, by introducing a tilt angle (*α, β*) in the *X* and *Y* directions between the reference and the object beams. The electric fields incoming from the reference and the object arms can therefore be described as Ψ_*R*_ = *P*_*R*_ and Ψ_*O,α*_ = Ψ_*O*_*e*^*i*(*αx*+*βy*)^ = *P*_*O*_*e*^*i*(Δ*ϕ*+*αx*+*βy*)^ respectively, where Ψ_*O,α*_ is the electric field Ψ_*O*_ with the added tilt, and *P*_*R*_ and *P*_*O*_ are real amplitudes. The electric fields combine on the image plane to form the interferogram

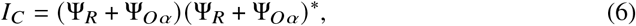

where ^*^ signifies the complex conjugate operation. Under the assumption that *P*_*R*_ is constant, *I*_*C*_ is described in the Fourier domain as

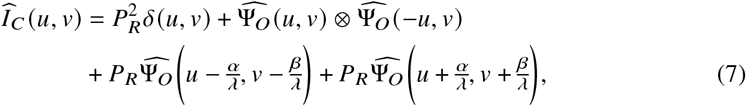

where *δ*(*u, v*) is a Dirac delta function and 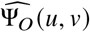 is the Fourier transform of Ψ_*O*_. Note that 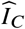 contains two sidelobes, which are represented in Eq. 7 by the terms 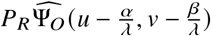 and 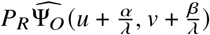. A graphical representation of both *I*_*C*_ and its Fourier transform can be found on the top left and right panels of Fig. 3, respectively.

**Fig. 2.**
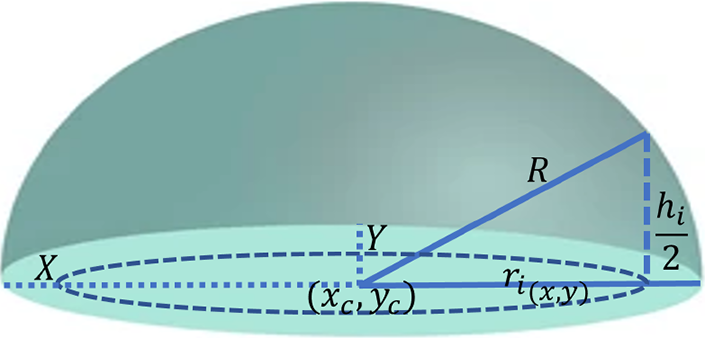
Schematic diagram for the estimation of sample thickness at each pixel position. Each pixel in the hemisphere of radius R has a height of *h*/2 and is located at a position *r* from the centre of the circular plane.

**Fig. 3.**
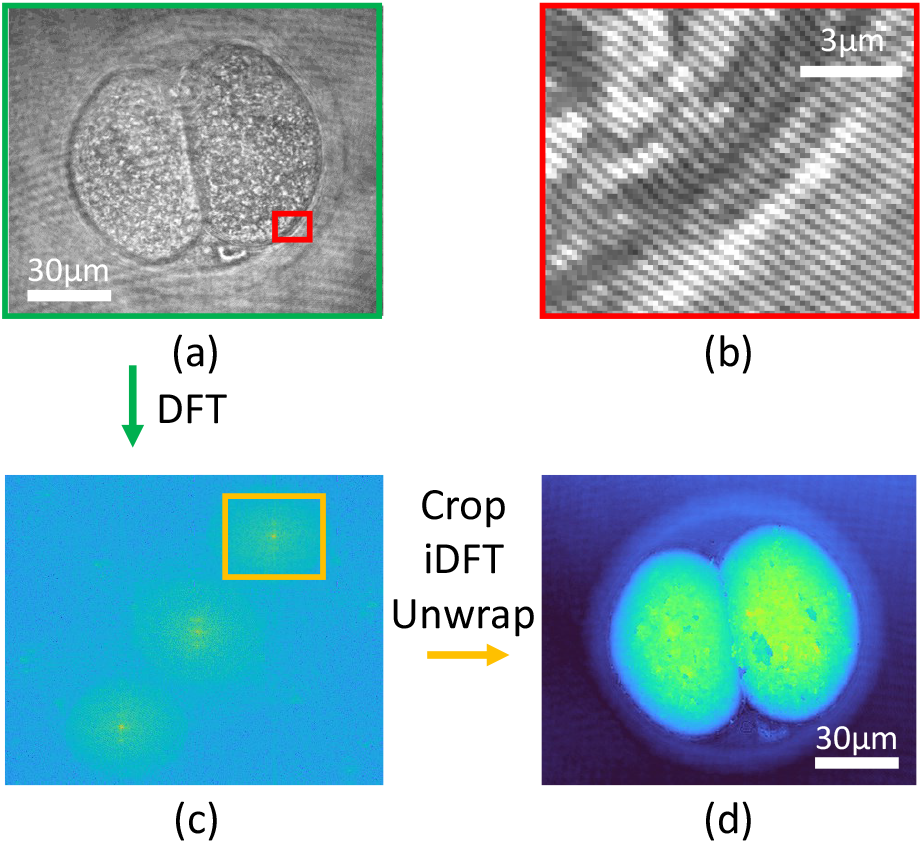
Flow diagram of the demodulation process, showing actual results from DHM images. The interferogram as detected by the CCD (*a*) contains modulated fringes that can be seen in the zoomed in panel (*b*). The demodulation starts by computing the discrete Fourier transform (DFT). On the Fourier plane (*c*), a sideband is cropped, as shown by an orange rectangle and an inverse DFT is used to retrieve an estimation of the electric field at the object plane. Panel (*d*) shows the estimated phase difference Δ*ϕ*, after unwrapping.

Due to the tilt angle (*α, β*), these sidelobes do not overlap with the other terms and can therefore be demodulated to retrieve Ψ_*O*_. This is done by computing the Fourier transform of *I*_*C*_, cropping and centering one of the sidelobes and then performing an inverse Fourier transform. This process, which retrieves *P*_*R*_Ψ_*O*_, is represented in Fig. 3

The last steps are to retrieve the angle of the electric field and then to unwrap it to avoid artificial discontinuities. This is done using the Herraez method which is based on unwrapping points with higher reliability values rather than on a continuous path [22]. This provides a faster reconstruction as well as consistent results when discontinuities or noisy areas are present.

### 2.6. Statistical analysis

All phase retrievals and RI index calculations were carried out in MATLAB version 2020b, while statistical analyses were performed using GraphPad Prism version 9 for Windows 10 (GraphPad Holdings LLC, CA, USA). Data were checked for normality and subjected to appropriate statistical testing as described in the figure legends. Statistical significant differences were set at P<0.05.

## 3. Results and discussion

Figure 4 (a and b) shows bright-field images acquired using the DHM setup when 2-cell stage embryos were cultured in low- or high-lipid containing growth media, respectively. We found that the phase information profiles extracted from the holograms of these bright-field images were vastly different for 2-cell stage embryos cultured in low-lipid media (Fig. 4c) compared to those cultured in high-lipid media (Fig. 4d), despite the similarity in overall morphology (Fig. 4a and b). Notably, the phase shift in the centre of each cell in the high lipid group was higher than for the low lipid group (Fig. 4d vs. 4c). This is clearly visualised in the reconstructed 3D topographical profiles of these embryos (Fig. 4f and 4e). Following that, we extracted phase information for each pixel within the region of interest (ROI) to compute the corresponding refractive index from Eq. 5. These RI profiles were calculated within the ROI with the background information subtracted. The ROI encompassed the area within and including the zona pellucida (protein-coat that surrounds the embryo), with the radius of the ROI being *R* in Eq.5. Representative 2D refractive index profiles are shown in Fig. 4g for low-lipid and Fig. 4h for high-lipid group with the corresponding 3D topographical RI profiles shown in Fig. 4i vs. 4j respectively. A comparison of the histogram distributions of RI between Fig 4g vs 4h is plotted in Fig 4k. Each histogram consists of a double peak where the low index peak (left) shows the RI of the zona pellucida, and fluid between the cells and the zona, while the high index peak (right) shows the RI within the cells of the embryo. Summarising these observations, while the RI in the zona regions were similar, an increased RI was observed within the cells of embryos that were cultured in high-lipid compared to those cultured in low-lipid media. This is in contrast to visual inspection (bright-field images) where no observable difference could be discerned between embryos cultured in the low- vs high-lipid containing media. This confirms that our DHM approach is capable of detecting small changes in lipid abundance within the embryo not visible to the naked eye. Our RI results are comparable to results obtained with a commercial optical diffraction tomography (ODT) system [11]. The advantage in the use of DHM is that the 3D RI profiles can be obtained from a single-shot hologram as opposed to reconstruction from multiple 2D holograms recorded at various incident angles as used in ODT.

**Fig. 4.**
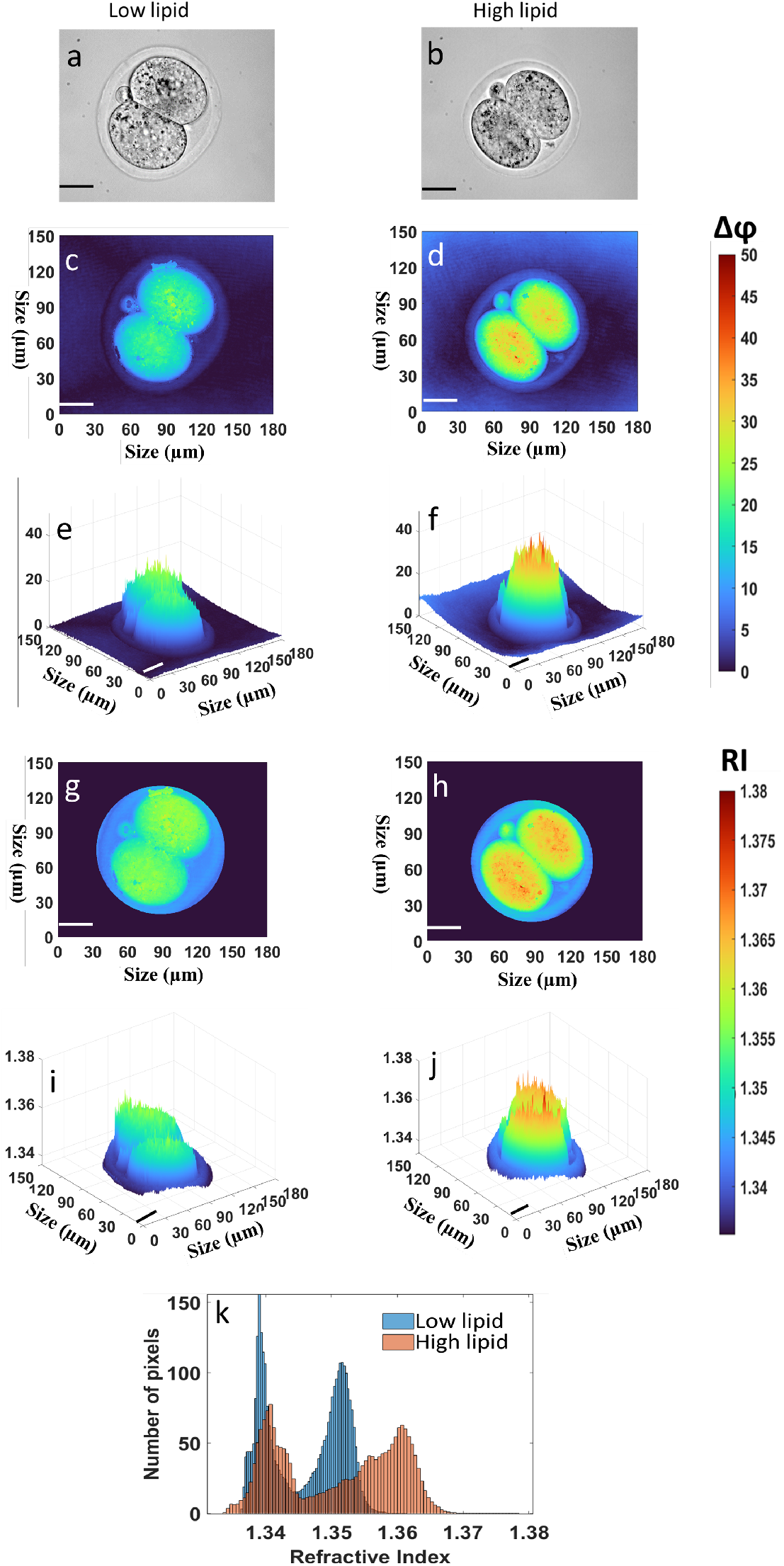
Representative phase information and refractive index maps for 2-cell stage embryos cultured in low- and high-lipid media. (a,b) Bright-field images of 2-cell stage embryos cultured in media containing low or high levels of lipid, respectively. (c,d and e,f) represents the reconstructed 2D phase profiles and their corresponding 3D topographical profiles for the low-lipid and high-lipid 2-cell stage embryo groups, respectively. (g,h) represents the reconstructed 2D refractive index profiles for the low- and high-lipid groups respectively with (i,j) their corresponding 3D topographical refractive index profiles within the ROI after background subtraction. (k) Frequency distribution of all pixels measured using the reconstructed 2D refractive index profiles of embryos cultured in low- and high-lipid containing media in (g) and (h), respectively. Scale bars = 30 μm.

To characterize dynamic changes in RI during pre-implantation development, embryos were cultured in either standard conditions (Low lipid; first and second column) or within media containing high lipid (third and fourth column). Embryos were imaged at the 4-cell (Fig. 5; first row), 8-cell (second row), morula (third row), and blastocyst-stage (fourth row). The phase profile and RI map for each developmental stage are shown in Supplementary Figures S1-S4. Within the low- and high-lipid groups, the highest RI was observed at the morula-stage (Fig. 5; third row). Visually, embryos cultured in high-lipid medium had an elevated RI at all stages of development with the exception of the blastocyst-stage (Fig. 5). This observation was confirmed following quantification of RI for each pixel. There was a consistent right-skewed second peak demonstrating a higher RI when embryos were cultured in high-lipid media throughout development, with the exception of the blastocyst-stage (Fig.5; fifth column). Following this, we quantified the average RI in multiple embryos for each developmental stage (see Fig 6a-e and Supplementary Table S2). The results were concordant with the representative data in Fig. 5.

**Fig. 5.**
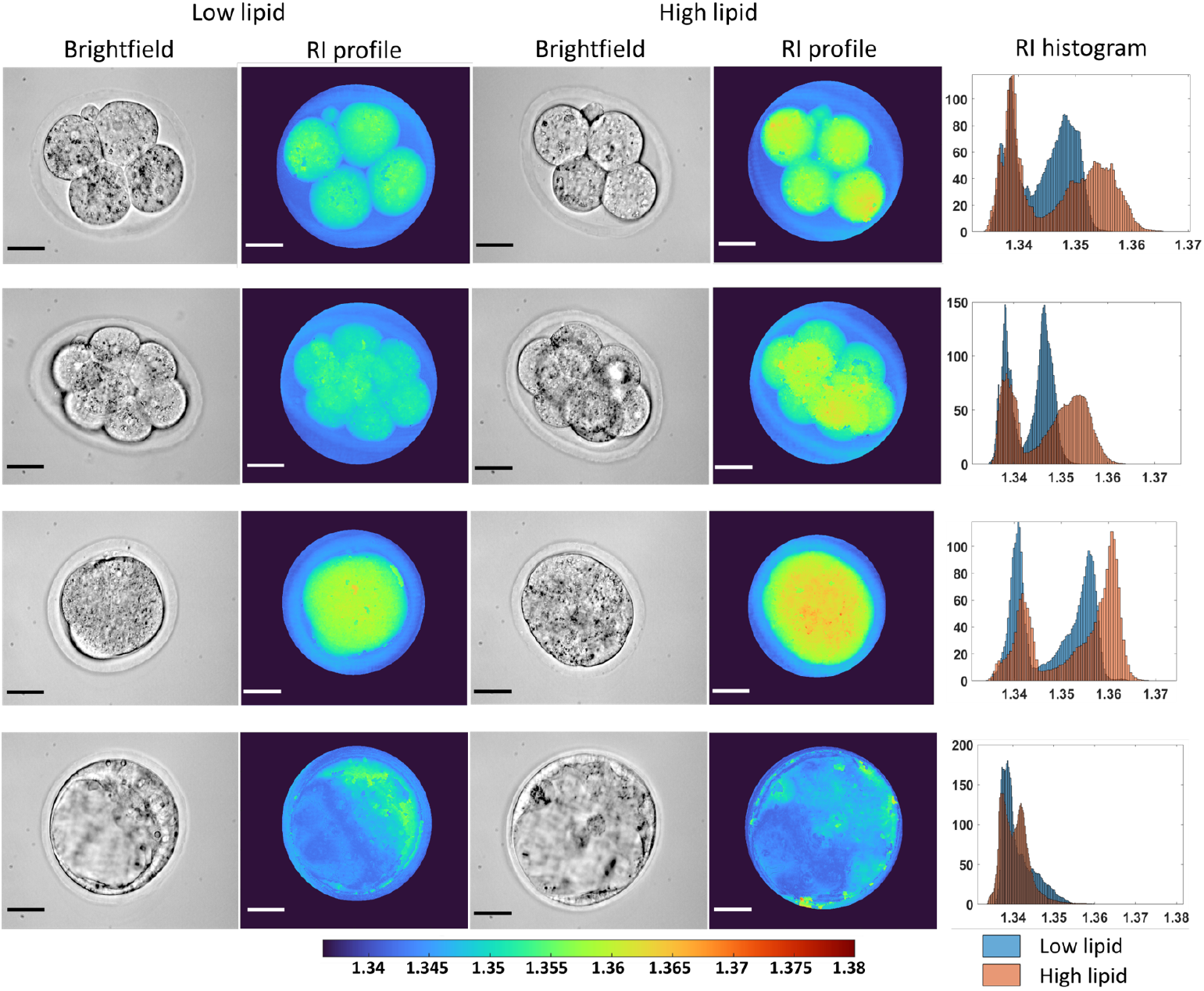
Representative refractive index profiles for 4-cell, 8-cell, morula and blastocyst stage embryos cultured in low- and high-lipid media. Rows 1-4 represent 4-cell, 8-cell, morula and blastocyst-stage embryos respectively. Columns 1 and 3 show the bright-field images with the corresponding RI profiles shown in columns 2 and 4 for low- and high-lipid embryos, respectively. Column 5 displays a comparison of the histogram distribution of RI recorded for each pixel between low- and high-lipid embryos at each stages of development. Scale bars = 30 μm.

**Fig. 6.**
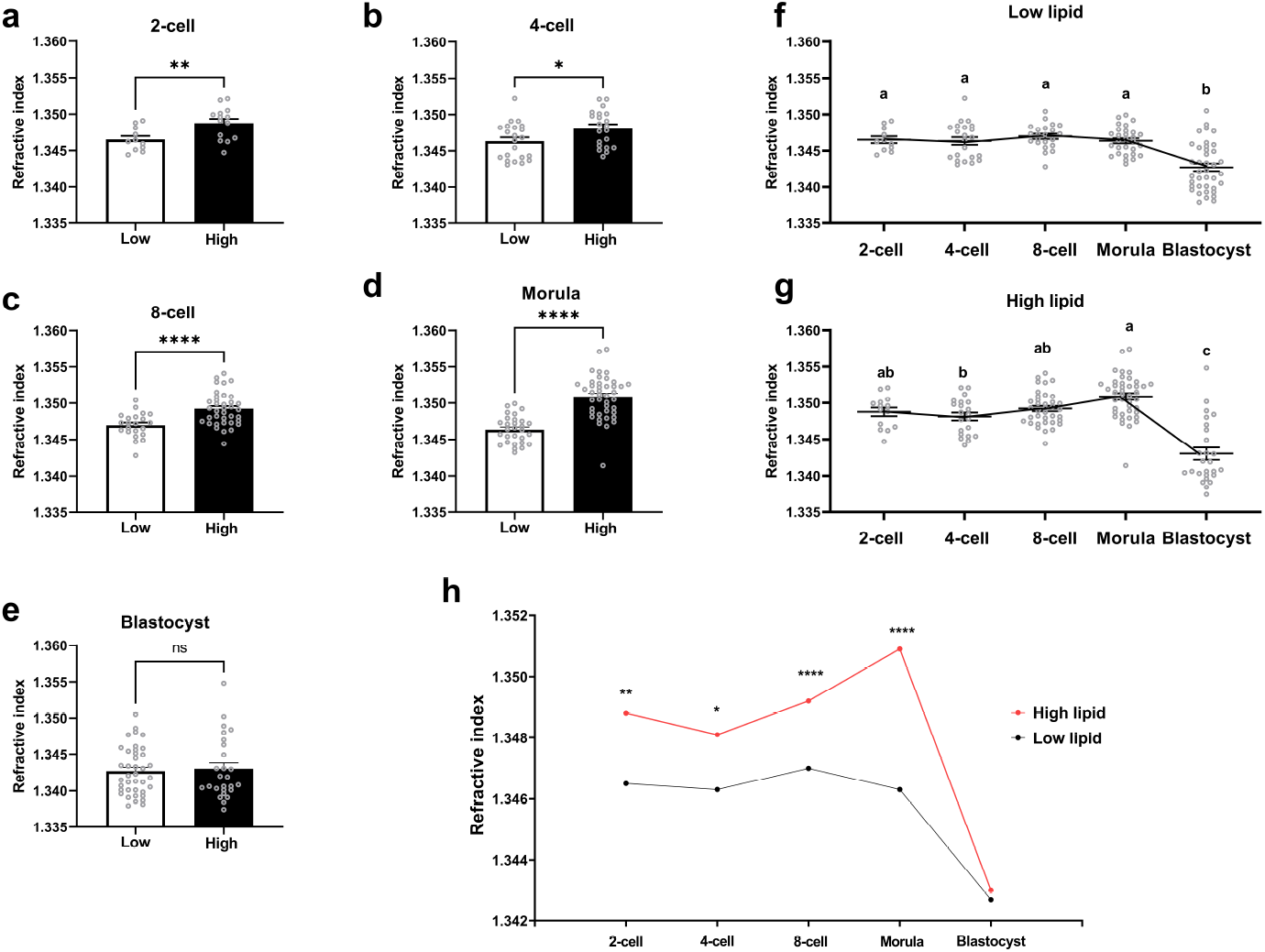
Refractive index throughout developmental stages significantly increased for embryos cultured in high-lipid media. The refractive index for all embryos was computed from their corresponding phase information profiles at the (a) 2-cell, (b) 4-cell, (c) 8-cell, (d) morula and (e) blastocyst-stage. Changes in refractive index throughout embryo development were recorded within the (f) low-lipid and (g) high-lipid groups of embryos. The refractive index for all embryos throughout development was also presented collectively to illustrate the relationship between the two treatment groups (h; *low- vs. high-lipid*). Data are presented as mean ± SEM, from 3 independent experimental replicates. Data were analyzed using either a Mann-Whitney test (a), an unpaired t-test (b,c,d,e,h) or one-way ANOVA with Tukey’s multiple comparison test (f,g). Asterisks indicate statistical significance between treatment groups within a particular developmental stage (*low- vs. high-lipid*), while different superscripts indicate statistical significance between developmental stages (*2-cell vs. 4-cell vs. 8-cell vs. Morula vs. Blastocyst*). *P<0.05 **P<0.01 ****P<0.0001

Interestingly, within both low lipid and high lipid groups, the RI remains relatively stable throughout development until the blastocyst-stage when it dramatically and significantly decreased (Fig. 6f and g; and presented collectively in 6h). A plausible explanation for this sharp reduction in RI is that it is due to the volumetric expansion of the blastocoel (fluid-filled) cavity as the embryo develops from a densely packed cluster of cells at the morula-stage to an embryo with a fluid-filled centre at the blastocyst-stage. This fluid-filled cavity may affect how light traverses through the embryo and therefore the recorded refractive index. Nonetheless, in support of our results, the RI recorded are within the range of indices previously reported for porcine embryos [9] and other cell types [23].

To demonstrate that the increase in RI was a result of elevated intracellular lipid, we quantified the abundance of lipid within embryos at each developmental stage qualitatively. In a separate cohort, embryos were again cultured in low- or high-lipid containing media and intracellular lipid quantified using a lipid-specific stain, BODIPY. With the exception of the 2-cell stage embryo. embryos cultured in high-lipid media contained a significantly higher abundance of intracellular lipid (see Fig 7). We postulate that the null difference observed for 2-cell stage embryos may be due to the relatively short duration in which they spent in high-lipid medium (6 h as opposed to all other stages, > 10 h). Despite the short period of culture within high-lipid medium, DHM was able to detect a shift in RI (see Fig. 4 and 6a). This indicates that DHM is more sensitive in detecting changes in lipid abundance compared to BODIPY staining.

**Fig. 7.**
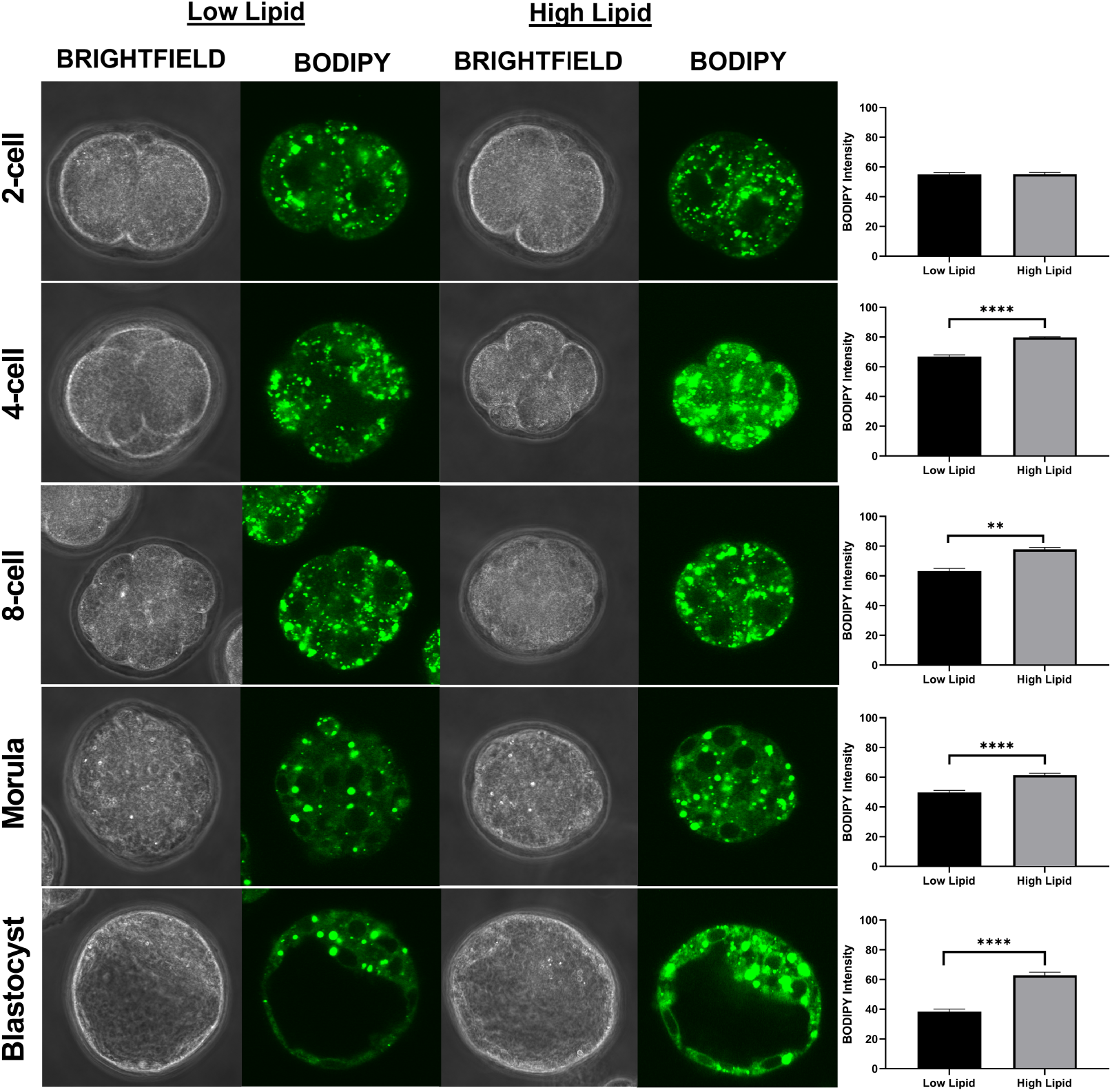
Embryos cultured in media containing high levels of lipid significantly increased lipid abundance within resultant embryos. Two-cell stage embryos were either cultured in media containing low (first and second column) or high levels of lipid (third and fourth column) and allowed to develop until the blastocyst-stage. Lipid abundance in resultant embryos at each developmental stage was quantified using the fluorescence intensity of BODIPY 493/503 from the maximum intensity z-projection generated for each embryo. Data are presented as mean ± SEM. n = 25-30 embryos per developmental stage, from 3 independent experimental replicates. Images were captured at 60x magnification. **P<0.01 ****P<0.0001

In contrast, lipid abundance in blastocyst-stage embryos was significantly higher following culture in high-lipid media compared to those cultured in low-lipid containing media (Fig. 7; fifth row). However, DHM did not detect a difference between blastocyst-stage embryos from the two groups (see Fig. 5; fourth row and 6e). We contend that this is likely due to the vast majority of contribution to the RI measurement for a blastocyst-stage embryo can be attributed to the fluid-filled cavity that develops at this stage.

A limitation of this present study is that our computed RI for the blastocyst-stage embryos was influenced by the fluid-filled centre. This highlights the potential consideration for future studies to focus on specific regions of the embryo, for example, limiting the region of interest to the cell-rich region of the blastocyst, termed the inner cell mass, which goes on to form the fetus (see Fig. 7; fifth row). As a result, DHM may prove most powerful in predicting embryo quality by imaging at earlier stages of development prior to development of the blastocoel cavity. It is noteworthty that a previous study has shown a clear linear relationship between the fluorescence quantum yield of BODIPY and refractive index [24]. Here, establishing a direct correlation between intracellular lipid levels and refractive index was outside of the scope for this study as separate cohorts of embryos were used for DHM imaging and the quantification of lipid abundance by immunohistostaining. This will be the subject of future studies. Further, it would also be of interest to quantify the absolute amount of lipids using alternate methods (such as mass spectrometry) to elucidate the relationship between intracellular lipids and refractive index. It would also be of interest to include transferring embryos that were cultured in the presence of serum into pseudopregnant female mice following DHM imaging to assess for potential long-term developmental anomalies typically associated with embryo overgrowth. Therein lies an opportunity where DHM imaging may act as a predictor of postnatal outcomes and fetal health.

## 4. Conclusion

To the best of our knowledge, this is the first reported use of DHM to characterize spatio-temporal changes in refractive index throughout pre-implantation embryo development. Importantly, this optical approach uses very low (micro-watt) excitation powers for short time durations, and therefore it is considered safe in terms of phototoxicity. The significance of this work lies in the capacity to rapidly and non-invasively acquire intrinsic phase change information from a developing cell. In this study, this refers to the changes in lipid content, and thus of refractive index within an embryo throughout development. The label-free nature of DHM also makes it ideal to form part of a multimodal analysis when combined with other optical techniques such as Raman spectroscopy [25, 26], hyperspectral imaging or FLIM. This work represents a step forward for DHM imaging to be part of an optical assessment of embryos in the IVF clinic to augment conventional visual assessment for embryo quality.

## 5. Acknowledgments

The authors would like to thank Dr. Graham Bruce for the critical reading of the manuscript and constructive suggestions. We acknowledge Dr. Philip Wijesinghe and Dr. Morgan Facchin for their useful discussions.

## 1. FIGURES AND TABLES

**Table S1.**
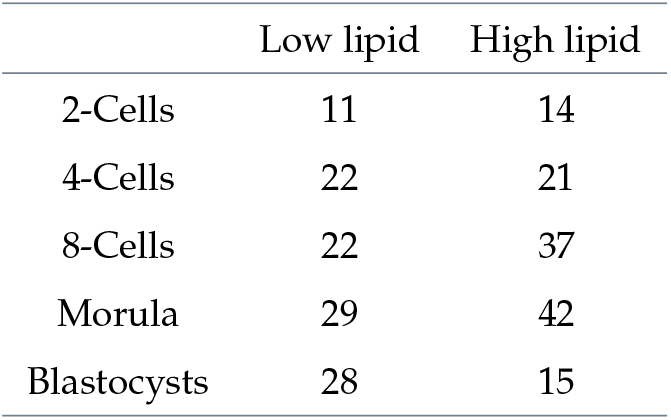
Sample sizes for the different embryo developmental stages imaged

|  | Low lipid | High lipid |
| --- | --- | --- |
| 2-Cells | 11 | 14 |
| 4-Cells | 22 | 21 |
| 8-Cells | 22 | 37 |
| Morula | 29 | 42 |
| Blastocysts | 28 | 15 |

**Table S2.** Average measured refractive indices for each developmental stage

|  | 2-cell | 4-cell | 8-cell | Morula | Blastocysts |
| --- | --- | --- | --- | --- | --- |
| Low lipid | 1.347 | 1.346 | 1.347 | 1.346 | 1.344 |
| High lipid | 1.349 | 1.348 | 1.349 | 1.351 | 1.346 |

**Fig. S1.**
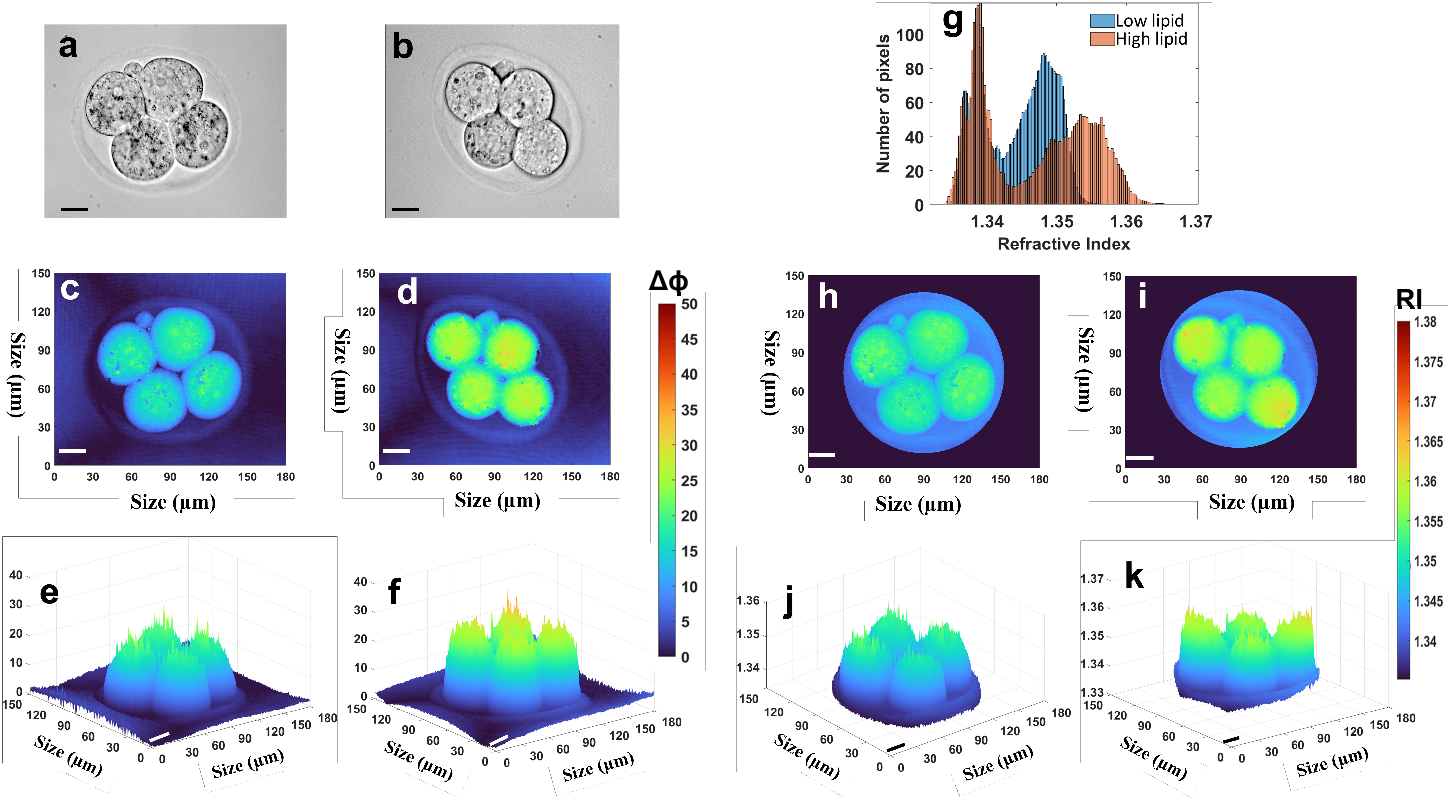
Representative phase information and refractive index maps for 4-cell embryos cultured in low or high lipid media. (a,b) Bright field images for low lipid and high lipid-treated 4-cell embryos, respectively. (c,e and d,f) Reconstructed 2D phase profiles and their corresponding 3D topographical profiles for low lipid and high-lipid group respectively. (h,j and i,k) Reconstructed 2D refractive index profiles for low lipid and high lipid group with their corresponding topographical refractive index profiles. (g) Frequency distribution of all pixels measured using the reconstructed 2D refractive index profiles for low and high lipid groups of embryos in (h and i, respectively). Scale bars = 20 *µ*m.

**Fig. S2.**
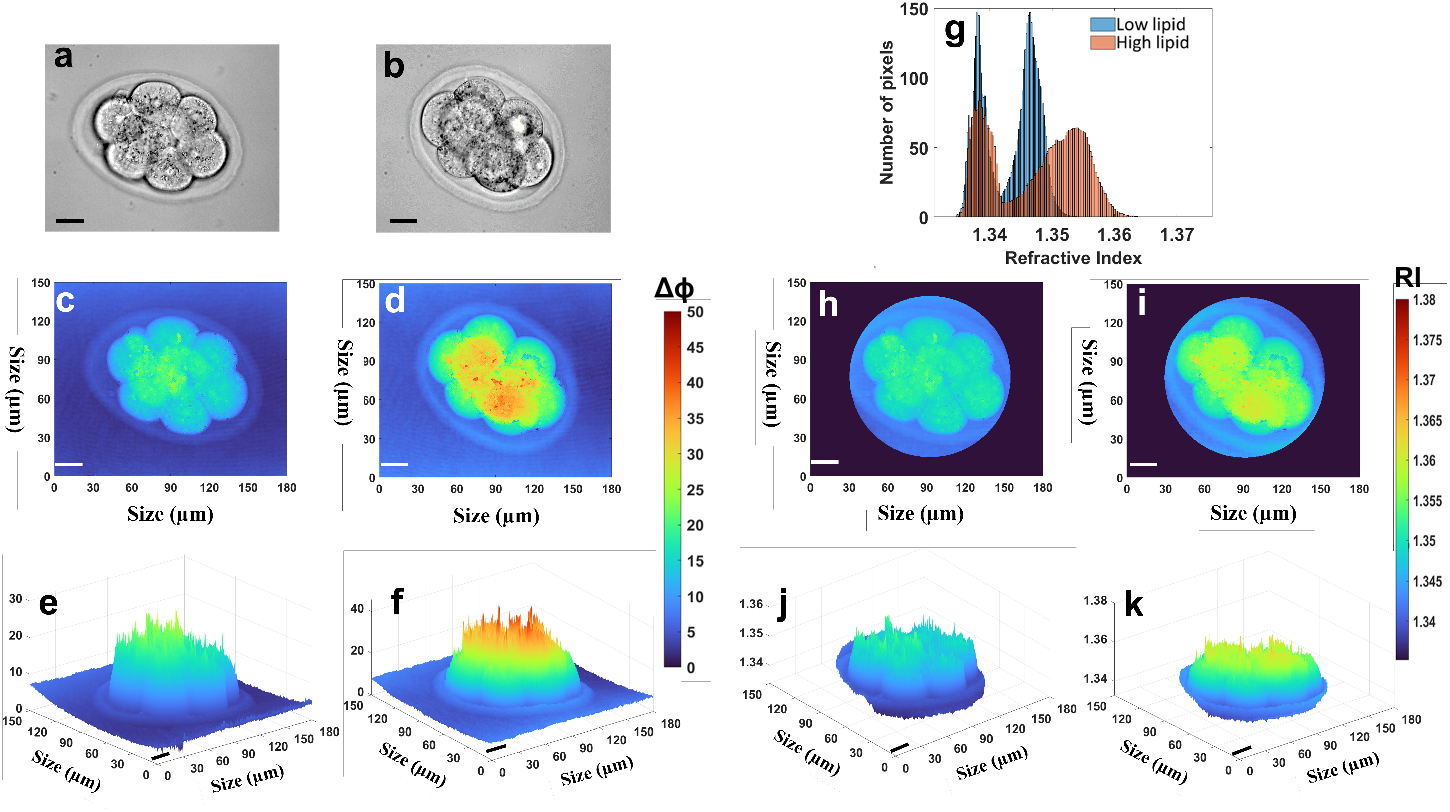
Representative phase information and refractive index maps for 8-cell embryos cultured in low or high lipid media. (a,b) Bright field images for low lipid and high lipid-treated 8-cell embryos, respectively. (c,e and d,f) Reconstructed 2D phase profiles and their corresponding 3D topographical profiles for low lipid and high-lipid group respectively. (h,j and i,k) Reconstructed 2D refractive index profiles for low lipid and high lipid group with their corresponding topographical refractive index profiles. (g) Frequency distribution of all pixels measured using the reconstructed 2D refractive index profiles for low and high lipid groups of embryos in (h and i, respectively). Scale bars = 20 *µ*m.

**Fig. S3.**
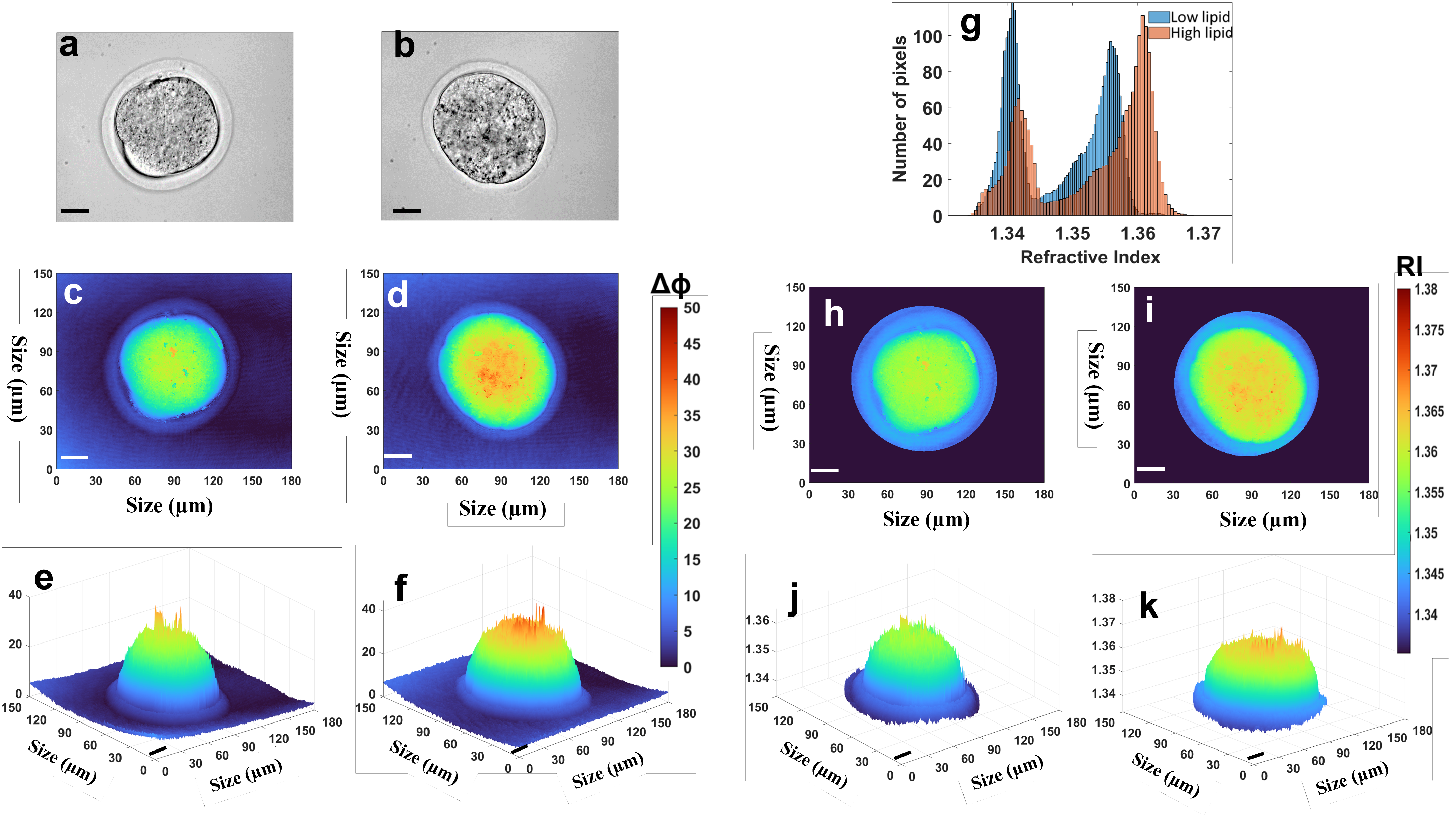
Representative phase information and refractive index maps for morula-stage embryos cultured in low or high lipid media. (a,b) Bright field images for low lipid and high lipid-treated morula-stage embryos, respectively. (c,e and d,f) Reconstructed 2D phase profiles and their corresponding 3D topographical profiles for low lipid and high-lipid group respectively. (h,j and i,k) Reconstructed 2D refractive index profiles for low lipid and high lipid group with their corresponding topographical refractive index profiles. (g) Frequency distribution of all pixels measured using the reconstructed 2D refractive index profiles for low and high lipid groups of embryos in (h and i, respectively). Scale bars = 20 *µ*m.

**Fig. S4.**
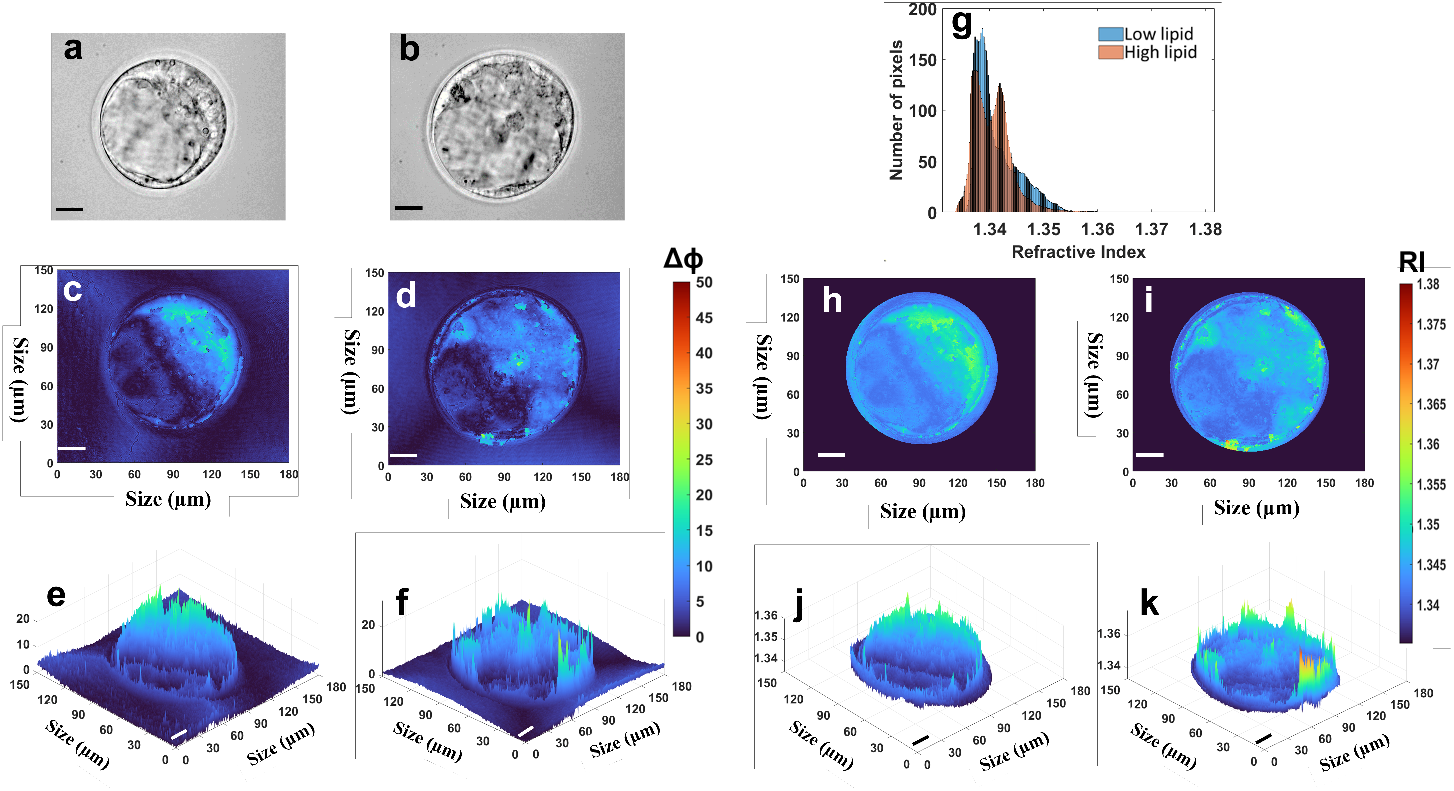
Representative phase information and refractive index maps for blastocyst-stage embryos cultured in low or high lipid media. (a,b) Bright field images for low lipid and high lipid-treated blastocyst-stage embryos, respectively. (c,e and d,f) Reconstructed 2D phase profiles and their corresponding 3D topographical profiles for low lipid and high-lipid group respectively. (h,j and i,k) Reconstructed 2D refractive index profiles for low lipid and high lipid group with their corresponding topographical refractive index profiles. (g) Frequency distribution of all pixels measured using the reconstructed 2D refractive index profiles for low and high lipid groups of embryos in (h and i, respectively). Scale bars = 20 *µ*m.

